# Coding *de novo* mutations identified by WGS reveal novel orofacial cleft genes

**DOI:** 10.1101/2020.04.01.019927

**Authors:** Madison R. Bishop, Kimberly Diaz Perez, Miranda Sun, Samantha Ho, Pankaj Chopra, Nandita Mukhopadhyay, Jacqueline B. Hetmanski, Margaret A. Taub, Lina M. Moreno-Uribe, Luz Consuelo Valencia-Ramirez, Claudia P. Restrepo Muñeton, George Wehby, Jacqueline T. Hecht, Frederic Deleyiannis, Seth M. Weinberg, Yah Huei Wu-Chou, Philip K. Chen, Harrison Brand, Michael P. Epstein, Ingo Ruczinski, Jeffrey C. Murray, Terri H. Beaty, Eleanor Feingold, Robert J. Lipinski, David J. Cutler, Mary L. Marazita, Elizabeth J. Leslie

## Abstract

While *de novo* mutations (DNMs) are known to increase risk of congenital defects, DNMs have not been fully explored regarding orofacial clefts (OFCs), one of the most common human birth defects. Therefore, whole-genome sequencing of 756 case-parent trios of European, Colombian, and Taiwanese ancestry was performed to determine the contributions of coding DNMs to OFC risk. Overall, we identified a significant excess of loss-of-function DNMs in genes highly expressed in craniofacial tissues, as well as genes associated with known autosomal dominant OFC syndromes. This analysis also revealed roles for zinc-finger homeobox domain and SOX2-interacting genes in OFC etiology.

## Introduction

Orofacial clefts (OFCs) are the most common craniofacial malformation in humans, affecting 1 in 1000 live births around the world^1^, and cause a significant personal, financial, and societal burden^2^. OFCs are phenotypically and etiologically heterogeneous which presents important opportunities and significant challenges for discovering causal genes. Phenotypically, these malformations can be divided into three general categories: clefting of the lip only (CL), clefting of both the lip and the palate (CLP), and clefting of the palate only (CP). Since CL and CLP (together denoted CL/P) share a defect in the primary palate, they are often analyzed together, while CP are typically analyzed separately because the secondary palate forms after the lip during development^2^. Further, OFCs can also be classified as an isolated (e.g. nonsyndromic) occurrence comprising approximately 70% of CL/P and 50% of CP cases or as a part of a syndrome comprising the remaining 30% and 50%, respectively.

These phenotypic classifications inform hypotheses about the type and number of genetic variants controlling risk of OFC. For example, syndromic OFCs are typically attributed to single gene mutations, chromosomal anomalies, or teratogens whereas nonsyndromic OFCs are considered to have a complex etiology with multiple genetic and environmental risk factors. Similarly, 15% of OFC cases have some family history of OFC, suggesting a major contribution of inherited genetic factors while the sporadic nature of most OFCs suggests a possible role for *de novo* mutations (DNMs). Of course, genetic variants of all types and frequencies plus environmental risk factors combine to influence the penetrance and expression of genes controlling OFCs, making these categories useful but not absolute.

Multiple large-scale genome-wide association studies (GWAS) have been performed in nonsyndromic OFC cohorts to identify ∼45 genetic loci which collectively account for ∼25% of the estimated liability to OFCs^3^. Exome and genome sequencing have been applied to identify genetic causes of syndromic OFCs in some family-based studies of nonsyndromic OFCs, but these approaches have not yet been applied to investigate primarily rare variants in a large number of samples. In this manuscript we focus on DNMs as one understudied class of variants influencing risk to OFCs. DNMs arise spontaneously in either the germ cell of the parents or during the early stages of embryonic development^4^. On average, individuals have about 100 DNMs throughout their entire genome, with approximately one DNM affecting the coding-region (exome)^51; 6^. Genetic studies of DNMs have led to previous successful identification of genes and pathways underlying multiple congenital disorders, such as congenital heart defects^7-9^, Kabuki syndrome^10^, and autism^11^. However, only a few DNMs have been reported for OFC to date^12^, and their overall contribution to OFCs has not been thoroughly assessed in a large sample or on a genome or exome-wide scale.

Whole genome sequencing (WGS) of case-parent trios ascertained through several studies of nonsyndromic OFCs generated as part of the Gabriella Miller Kids First Pediatric Research Consortium now makes it feasible to investigate DNMs contributing towards OFCs on a genome-wide scale. Therefore, we analyzed the contribution of coding DNMs to various aspects of OFC risk in case-parent trios of European, Colombian, and Taiwanese ancestry.

## Methods

### Sample of case-parent trios

This study summarizes the initial findings on de novo variants in three samples of case-parent trios, one of 415 trios of European ancestry recruited from sites around the United States, Argentina, Turkey, Hungary and Spain; a second set of 275 trios from Medellin, Colombia; and a third set of 125 trios from Taiwan. In this study, the three samples are referred to as European, Colombian, and Taiwanese, respectively. Recruitment of participants and phenotypic assessments were done at regional treatment centers after review and approval by each site’s institutional review board (IRB) and the IRB of the affiliated US institutions. Among parents, 88.7% of European, 93.1% of Taiwanese, and 100% of the Colombian parents were unaffected. As most OFC cases are likely to have multifactorial etiology, we consider DNMs to be one of many genetic factors that could influence risk and they need not be limited to cases without any family history of OFC. Therefore, as in our prior work ^13^, the cleft status of the parents (Supplementary Table 1) was not considered in these analyses.

### Whole-genome sequencing and variant calling

The OFC trios were sequenced as part of the Gabriella Miller Kids First Pediatric Research Consortium (https://commonfund.nih.gov/kidsfirst/overview), established in 2015 with the aim of addressing gaps in the understanding of the genetic etiologies of structural birth defects, such as OFCs, and pediatric cancers. The McDonnell Genome Institute (MGI), Washington University School of Medicine in St. Louis carried out the whole genome sequencing (WGS) of the European samples which were subsequently aligned to hg38 and variant called by the GMKF’s Data Resource Center at Children’s Hospital of Philadelphia. Sequencing of the Colombian and Taiwanese samples was conducted at the Broad Institute and data were aligned to hg38 and called using GATK pipelines ^6; 14; 15^ at the Broad Institute (https://software.broadinstitute.org/gatk/best-practices/workflow). Details of the alignment and genotyping workflow used to harmonize these three datasets were recently published ^16^; briefly, all samples were realigned and recalled using a GATK pipeline at the GMKF Data Resource Center. Overall, the WGS data from these three studies were quite comparable. The average depth per sample for all sequenced individuals was 29.16.

### Quality control

The WGS data for the 415 European, 275 Latino, 125 Taiwanese case-parent trios was evaluated based on a variety of quality metrics. Individuals with an average read depth, Mendelian errors, and missingness outside of three standard deviations from the mean were removed. Additional individuals were removed based on Ts/Tv, exonic Ts/Tv, Silent/Replacement, and Heterozygotes/Homozygotes ratios that were less or greater than expected while allowing for somewhat lower ratios of Heterozygotes/Homozygotes ratios in the Colombian sample (which included trios drawn from a number of consanguineous pedigrees). Family relationships were confirmed with identity-by-descent analyses conducted in PLINK (version 1.90b53). X-chromosome heterozygosity was used to confirm the sex of all individuals. Finally, after assessing each individual’s quality metrics, only complete case-parent trios were retained, leaving 374 European, 267 Colombian, and 116 Taiwanese case-parent trios for analysis.

### Identification of de novo variants

Called variants were filtered for minor allele count (MAC) ≥ 1, genotype quality (GQ) ≥ 20, depth (DP) ≥ 10, variant quality scores (QUAL) ≥ 200, and quality by depth (QD) ≥ 3.0 using VCFtools (version 0.1.13) and GATK (version 3.8); additionally, only biallelic variant calls were included in this study. To generate a list of high confidence DNMs, variants were further filtered based on allele balance (AB). An AB filter ≥ 0.30 and ≤ 0.70 was used for variant calls in the offspring, and an AB filter < 0.05 was used for the corresponding variant calls in the offspring’s parents. Annotation of high confidence DNMs was completed using ANNOVAR (version 201707). One additional European case-parent trio was removed after this annotation because the offspring of European ancestry had 328 whole-genome DNMs, which was greater than three standard deviations from the observed mean. A Chi Square goodness-of-fit test was used to confirm that the observed number of coding DNMs for each proband followed the expected Poisson distribution (P=0.93), with the value of lambda being equal to the mean of the number of coding DNMs per proband. The number of DNMs genome-wide for each proband did appear to differ between populations (p<2.2 × 10^−16^; Supplemental Figure 7A); however, this is most likely due to sequencing artifacts or ancestry bias within databases used in the processing and filtering pipeline and not necessarily attributed to some unknown biological event. DNMs were then further filtered based on annotation, and only rare (MAF <0.1% in gnomAD v2.0.1), coding DNMs were kept for further analysis. Although there was a significant difference between populations genome-wide, there was no difference between the number of coding DNMs (P=0.12; Supplemental Figure 7B).

### Statistical analysis of de novo variants

The statistical analysis used to determine if there was any excess of coding DNMs by variant class throughout the entire exome and by variant class in individual genes was carried out using the DenovolyzerByClass and the DenovolyzerByGene function, respectively, in DenovolyzeR (0.2.0). This R package compares the observed number of DNMs to the expected number based on chance using a well-established model developed by Samocha *et al*. (2014) ^17^. Under the null model of no association between mutation class and disease status, the number of observed DNMs is expected to be Poisson^18^, with mean determined by the sequence of the genes in the exome and the fixed sample size. Thus, for each class of mutation, we have a single Poisson distributed observation, with “known” mean M, and therefore known variance M, and standard deviation sqrt(M). Thus, M here is fixed “known” constant. Under the alternate model, the number of observed mutations, A, is also Poisson, but A does not necessarily equal M. Here we plot A/M, together with the exact 95% confidence interval of A (the values of A above 2.5 percentile and below the 97.5 percentile divided by M) determined from the Poisson. The DenovolyzerByClass function paired with the includeGenes option in the DenovolyzeR program was used to test if more DNMs were observed than expected in a particular set of genes. Again, 95% confidence interval were displayed for the enrichment values, as in the autism study by Satterstrom *et al* ^*19*^. Because prevalence differs between males and females, the rate of observed DNMs is expected to differ between males and females for any mutation exerting the same effect on both sexes on the liability scale. When a rate difference was first observed in children with autism, the observation was dubbed the “female protective effect,” but this general phenomena is best thought of as simply a result of a mutation having a constant effect on the liability scale and differing prevalence between the sexes^20^. Because our female cases also have a higher rate of DNMs, and a lower overall prevalence, we estimate the effect size of each class of DNMs on a liability scale, and observed the difference in DNM rates.

### Gene set analyses

GSEA was carried out using using ToppFun available through the ToppGene Suite (https://toppgene.cchmc.org/enrichment.jsp); P-values were adjusted for multiple testing using Benjamini and Hochberg^21^, a correction method available within ToppFun. The top five most significant terms for each assessed category were then compared. Marker genes expressed in ectodermal and mesenchymal cell clusters were identified using single cell RNAseq and can be found in the supplementary information published in “The Molecular Anatomy of Mammalian Upper Lip and Primary Palate Fusion at Single Cell Resolution”^22^. Another set of functionally relevant genes was assembled using processed RNAseq data generated from human neural crest (HNC) cell samples (GSM1817212, GSM1817213, GSM1817214, GSM1817215, GSM1817216, and GSM1817217), which was downloaded from the Gene Expression Omnibus (https://www.ncbi.nlm.nih.gov/geo/query/acc.cgi?acc=GSE70751) ^23; 24^. The six cranial HNC cell samples were derived from iPSCs/ESCs from three human individuals, and RNA-sequencing was carried out of using NEBNext Multiplex Oligos for sequencing on an Illumna HiSeq 2000. The expression levels from the six samples were averaged and then ranked from highest expression to lowest expression. Genes with pLI scores ≥0.95 and in the top 20^th^ percentile for HNC expression were prioritized and tested for an excess of DNMs in our trios compared to what would be expected by chance. The clinically-relevant set of genes was constructed after a thorough literature search; known and suggested genes harboring mutations associated with OFC are summarized in Supplementary Table 4. The list of suggestive and significant loci use to identify DNMs in genes within ±500kb was generated using data generated from Carlson et al. (2019)^25^ and Yu et al. (2017)^26^.

### In situ hybridization

Studies involving mice were conducted in strict accordance with the recommendations in the ‘Guide for the Care and Use of Laboratory Animals’ of the National Institutes of Health. The protocol was approved by the University of Wisconsin-Madison School of Veterinary Medicine Institutional Animal Care and Use Committee (protocol number 13–081.0). C57BL/6J mice (Mus musculus) were purchased from The Jackson Laboratory. Timed pregnancies were established as described previously^27^. Embryos at embryonic day (E) 11 were dissected in PBS and fixed in 4% paraformaldehyde in PBS overnight followed by graded dehydration (1:3, 1:1, 3:1 v/v) into 100% methanol for storage. After rehydration, embryos were embedded in 4% agarose gel and 50 μm sections through the lambdoidal junction were made using a vibrating microtome. In situ hybridization was performed as previously described^28^. Gene-specific ISH riboprobe primers were designed using IDT PrimerQuest (http://www.idtdna.com/primerquest) and affixed with the T7 polymerase consensus sequence plus a 5-bp leader sequence (CGATGTTAATACGACTCACTATAGGG) to the reverse primer (Supplementary Table 5). Sections were imaged using a MicroPublisher 5.0 camera connected to an Olympus SZX-10. For each gene, representative images were selected from staining conducted on at least three sections from independent mouse embryos.

## Results

A set of high-confidence DNMs from WGS was generated from 756 complete case-parent trios; counts by cleft subtype, ethnicity, and sex are presented in Supplementary Table 1. The majority of the offspring had a CL or CLP; 58 European offspring had CP only. While 97% of cases are reported to have an isolated OFC, 3.0% reported other features (e.g. speech delay, hypertelorism, and intellectual disabilities). However, none of these trios have been diagnosed with a recognized genetic syndrome with molecular confirmation. Overall, 73,027 DNMs were identified genome-wide in the 756 case-parent trios with an average of 96.60 DNMs per proband (Figure 1A). Although we identified DNMs genome-wide, this initial analysis reported here focuses on rare, coding DNMs. After filtering for rare (MAF <0.1% in gnomAD v2.0.1) exonic and splicing variants, 862 coding DNMs in 808 genes were identified, averaging 1.14 DNMs per trio (Supplementary Table 2 and 3) and following the expected Poisson distribution (p=0.93) (Figure 1B)^18^.

**Figure 1.**
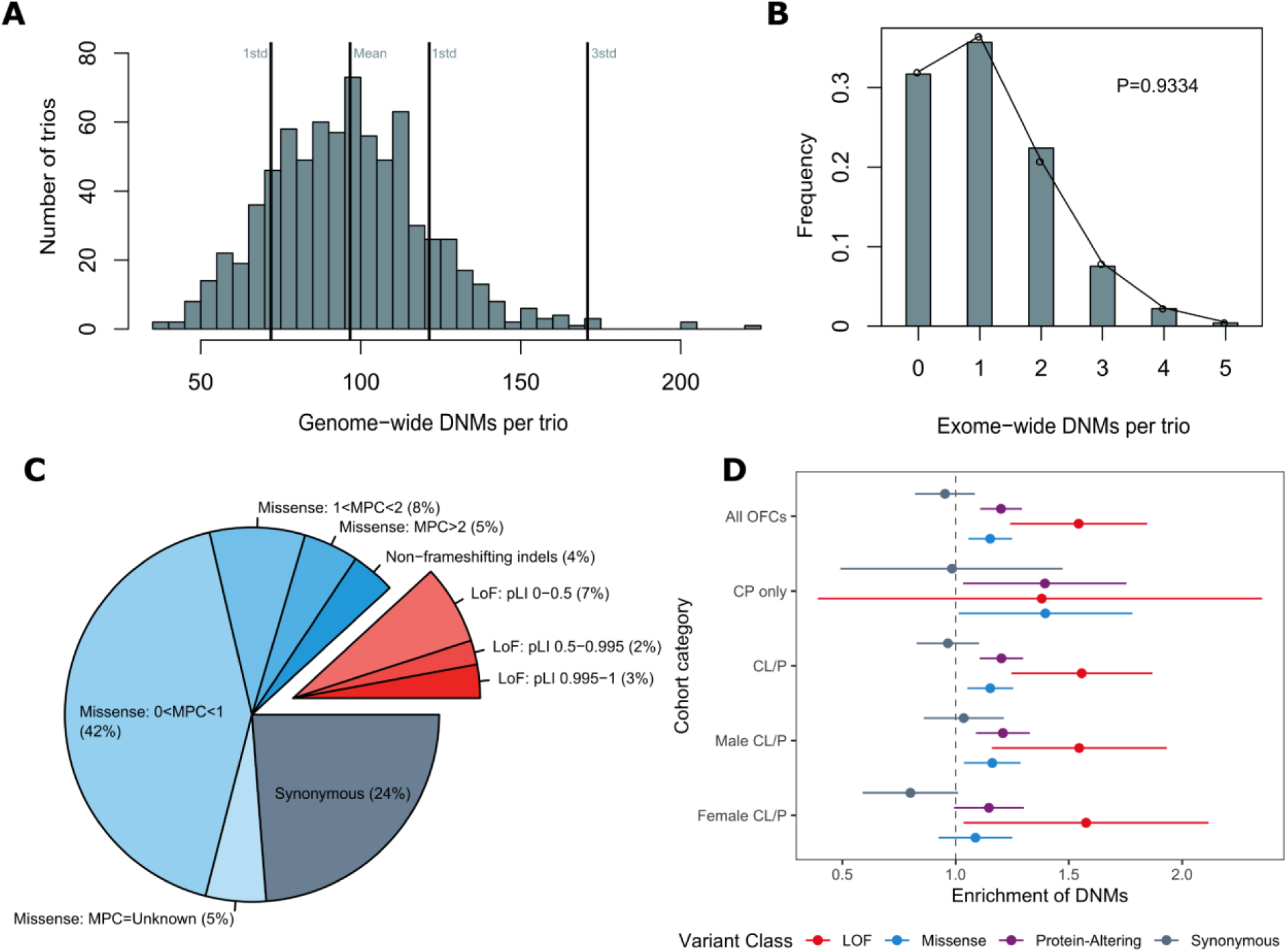
Loss of function and missense *de novo* mutations are enriched in cases with orofacial clefts. (A) Distribution of DNMs per trio genome-wide. (B) Distribution of coding DNMs per trio (C) Distribution of rare, coding DNMs by variant class for all OFC trios. DNMs were further divided by MPC score (missense) and gene pLI score (LoF) as a measure of potential deleteriousness. (D) Enrichment of DNMs ± two standard errors by variant class for all OFCs, probands with CP only, probands with CL/P, male probands with CL/P, and female probands with CL/P. The comparison of CP only trios by sex of the proband was not presented due to small sample sizes.

First, we characterized the distribution of types of coding DNMs amongst all trios and stratified by OFC subtypes and proband sex. We categorized coding DNMs into three classes based on their predicted function: missense (consisting of single amino acid changes and non-frameshifting insertions/deletions), predicted loss-of-function (LoF, made up of stop-gain, frameshifting insertions and deletions, and essential splice site variants), and a combined group of missense and LoF hereafter referred to as protein-altering DNMs. The majority of DNMs (64%) were missense variants (including 4% non-frameshifting indels), 12% were LoF, and the remaining 24% were synonymous (Figure 1C).

To assess the severity of specific types of DNMs, missense and LoF DNMs were additionally subcategorized by MPC score and pLI score respectively. MPC scores are a variant-specific combined measure of predicted deleteriousness that includes <u>M</u>issense badness, <u>P</u>olyPhen-2, and <u>C</u>onstraint. In contrast, the pLI scores are gene specific and represent the tolerance of a gene LoF variants; the more intolerant a gene is to a LoF variant the closer the pLI score is to 1. The overall proportion of DNMs in variant functional classes and subcategories were not significantly different for each OFC subtype (p=0.20; Supplementary Figure 1). The same was true for all OFCs when we compared the proportions of DNMs by sex of the proband (p=0.48; Supplementary Figure 2).

We next determined whether OFC trios possessed significantly more coding DNMs than expected based on mutational models^17^. The 756 trios had a significant excess of protein-altering DNMs (enrichment=1.20; *P*=3.22 × 10^−6^) (Figure 1D and Supplementary Table 2). The observed excess can be attributed to both LoF DNMs (enrichment=1.54; P=2.55 × 10^−5^) and missense DNMs (enrichment=1.15; P=5.92 × 10^−4^). As anticipated, trios did not possess an excess of synonymous DNMs (enrichment=0.95; *P* =0.76). When stratified by OFC subtypes, a significant excess of protein-altering DNMs was found among both CL/P trios (enrichment=1.20; *P*=6.12 × 10^−6^) and CP trios (enrichment=1.39; *P*=9.32 ×10^−3^), but the difference in strengths of association is difficult to interpret due to the differing sample sizes (Supplementary Table 2). Among CL/P females (n=254), the excess of protein-altering DNMs (enrichment=1.15; *P*=2.76 × 10^−2^) was primarily attributed to an excess of LoF DNMs (enrichment=1.58; *P*=7.28 × 10^−3^). In CL/P males (n=444), a similar excess for LoF DNMs (enrichment=1.55; *P*=9.69 × 10^−4^) and a significant excess of missense DNMs (enrichment=1.16, p=4.35 × 10^−3^) was observed. However, when the effects of DNMs in males and females with CL/P were compared directly on a liability scale, no significant differences were observed between male and female for any variant class (Supplementary Figure 3).

Next, a gene set enrichment analysis (GSEA) was performed to determine if genes with LoF or protein-altering DNMs clustered into specific gene sets or pathways relevant to craniofacial development. We also carried out GSEA for genes with synonymous DNMs as a control because these DNMs represent a set of variants most likely to have no effect on OFC risk. Protein-altering DNMs were enriched in genes belonging to gene sets broadly related to development and also more specific sets related to OFCs and craniofacial development (Supplementary Figure 4). For example, a significant enrichment of protein-altering DNMs was identified in genes related to the biological process term “embryo development” (*P*=6.11 × 10^−6^) and human disease terms such as “uranostaphyloschisis” (clefting) and “cleft palate” (*P*=3.28 × 10^−4^ and *P*=5.88 × 10^−4^, respectively). Similarly, an enrichment was identified for multiple terms that described abnormal embryo morphology in mouse (Supplementary Figure 4A, lower panel). Overall, we found that protein-altering DNMs were enriched in genes involved in embryonic development, craniofacial development, and human craniofacial disorders, whereas synonymous DNMs were not enriched in genes relevant to craniofacial development.

Because CL/P and CP have historically been considered distinct disorders, a GSEA was performed for genes with protein-altering DNMs in CL/P and CP separately to determine if this distinction was reflected in the WGS data. Overall, different gene ontology terms achieved statistical significance for offspring with CL/P and CP (Supplementary Figure 5). Many biological process and molecular function terms were statistically significant for offspring with CL/P whereas only one term (limb bud formation) was significant for offspring with CP only (*P*=6.32 × 10^−3^) (Supplementary Figure 5). For disease and human phenotype terms, “cleft palate”, “cleft palate (isolated)”, “congenital abnormality”, “hearing problem”, “osteogenesis imperfecta”, and “uranostaphyloschisis” were all significantly enriched for genes with protein-altering DNMs in the CP group but not for CL/P. In contrast, five mouse phenotypic terms were statistically significant in the CL/P trios, but only one mouse term (abnormal bone ossification) was significant for CP (*P*=3.37 × 10^−2^). Although both CL/P and CP genes were associated with terms related to embryonic development, the genes with DNMs in CL/P described broad embryonic development whereas the genes mutated in CP appeared more specific to the palate. The results of these analyses suggest that DNMs play an important role in both CL/P and CP and that these potential differences should be further investigated in a larger sample of CP only trios.

Although the GSEA results are promising, the significant terms represent very general problems with development. To get a more specific view of the impact of genes with DNMs in craniofacial development, we used recently published single-cell RNA sequencing data from the lambdoidal junction of the developing murine upper lip ^22^. The lambdoid junction is the point of fusion of three facial prominences creating the primary palate and upper lip, so we hypothesized that the marker genes for each cell cluster are among the best candidate genes in the genome to be involved in OFCs and could potentially harbor an excess of DNMs. To address this question, marker genes belonging to eight clusters of ectodermal cells and nine clusters of mesenchymal cells were analyzed (Figure 2). The marker genes from two ectodermal cell clusters (E5/E10 and E0/E11) positioned at the nasal process fusion zone and the olfactory epithelium had a significant excess of protein-altering DNMs (Figure 2A and 2C; *P*=1.38 × 10^−5^ and *P*=3.27 × 10^−5^, respectively). A single mesenchymal cell cluster, classified as Schwann cell progenitors, had a significant excess of protein-altering DNMs (Figure 2B and 2C;*P*=5.63 × 10^−4^); this cluster of cells was located adjacent to the fusing lambdoidal junction through mapping the mesenchymal clusters using *in situ* hybridization (Figure 2D) ^22^. Overall these analyses point to genes expressed in the cells at the point of fusion as being particularly relevant for risk of developing an OFC.

**Figure 2.**
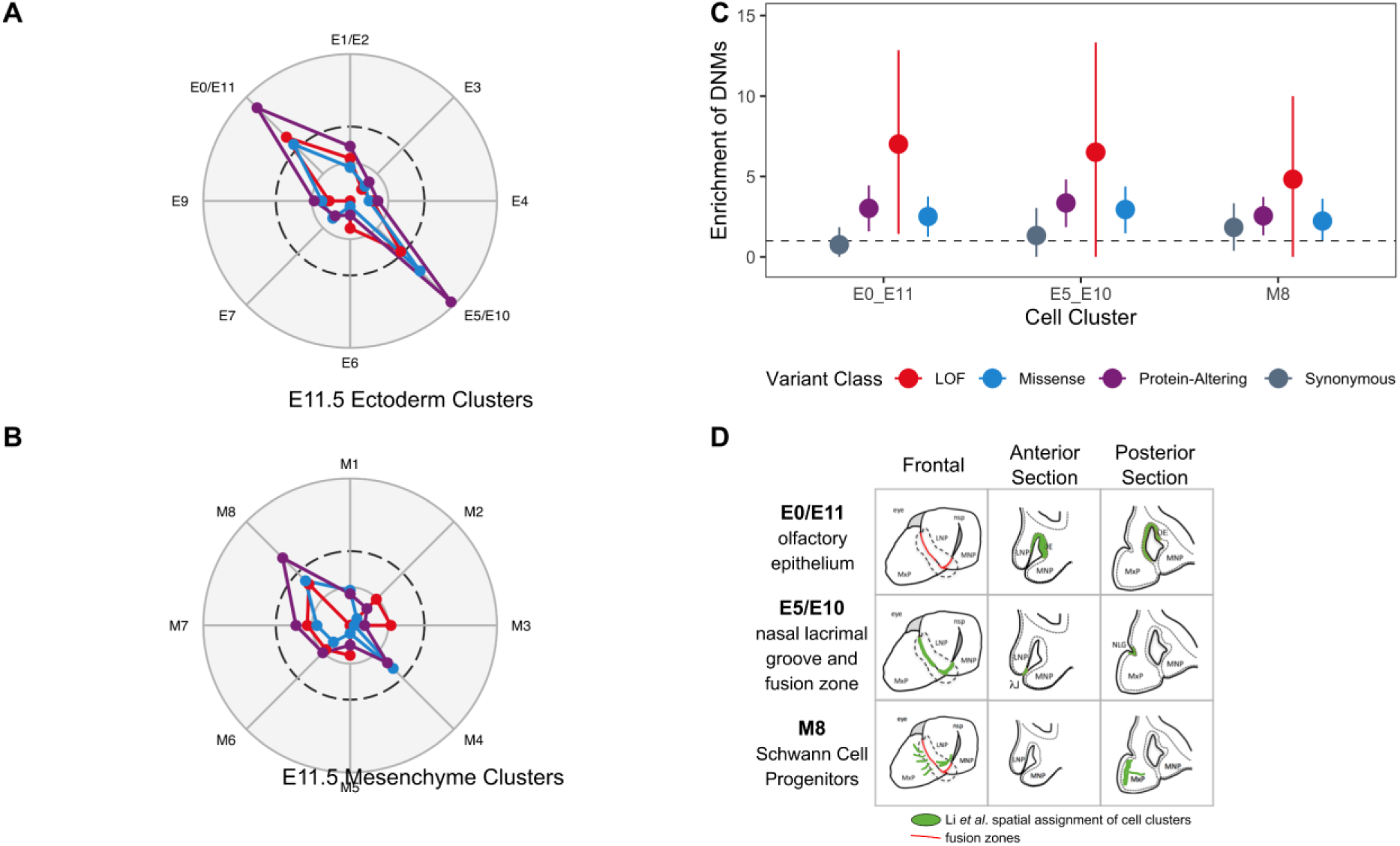
*De novo* mutations are enriched in genes expressed at the point of fusion in lip development. Marker genes for each ectodermal cell sub-cluster (A) and mesenchymal cell sub-cluster (B) were analyzed for an excess of DNMs, and the -log(p-value). In each radar plot, the dashed circle represents the significance threshold after correcting for multiple tests (p=2.9 × 10^−3^). The inner circle represents nominal significance (p=0.05); outer circle represents p=1.0 × 10^−5^. (C) Enrichment of DNMs ± two standard errors for significant cell sub-clusters. (D) Depiction of spatial assignment of cell clusters with significant excess of DNMs in the frontal view, anterior section, and posterior sections of the lambdoidal junction; adapted from Li *et al*.

We next identified individual genes with an excess of DNMs by comparing our observed mutation counts with published per-gene mutability models^4; 17^. Two genes (*TFAP2A* and *ZFHX4*) had significantly more LoF DNMs than expected after correcting for multiple testing (i.e. number of genes with any protein-altering DNM) (Figure 3). Notably, *TFAP2A* remained significant given a more conservative exome-wide significance threshold suggested by Ware *et al*. ^4^ (p < 1.3 ×10^−6^) despite only observing two distinct LoF DNMs (p.E104* and p.G145Efs*18; P=1.11 × 10^−6^). While multiple genes had more than one protein-altering DNM that may be critical to the underlying etiology of OFCs, only two genes (*TFAP2A* and *IRF6*) had a significant excess of protein-altering DNMs after correcting for multiple tests (Figure 3). In addition to the two LoF DNMs described above, we identified a third missense variant in *TFAP2A* (p.S247L). We also identified three missense variants in *IRF6* (p.G376V, p.N88D, and p.R84H).

**Figure 3.**
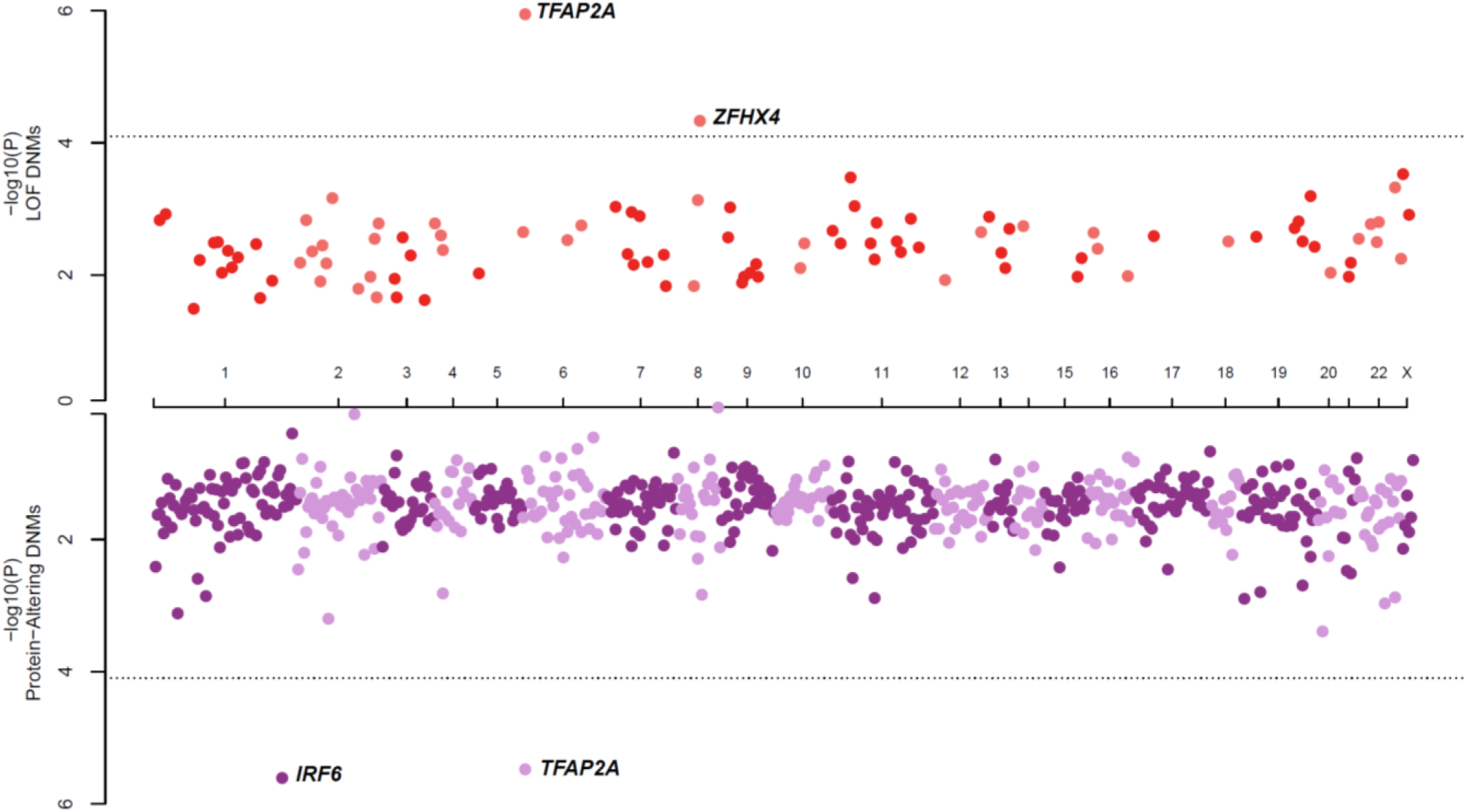
De novo mutations in *IRF6, TFAP2A*, and *ZFHX4* are associated with OFCs. Identification of single genes with an excess of LoF DNMs (top axis) or protein-altering DNMs (missense and/or LoF DNMs; bottom axis). The dashed line indicates the significance threshold after correcting for multiple tests is p<7.9×10^−5^.

We reasoned that genes highly expressed in craniofacial-relevant tissues that are intolerant to LoF variants would be good candidate genes for OFCs even if they do not individually reach formal statistical significance. To identify such genes, RNA-seq data from human neural crest cell lines (a cell type giving rise to a majority of facial structures) was used. A significant excess of LoF DNMs was observed among genes with pLI>0.95 in the top 20^th^ percentile of neural crest gene expression (enrichment = 2.49; P=3.46 × 10^−6^, Figure 4). The 132 genes with protein-altering DNMs in this category were significantly enriched for gene ontology terms related to gene regulation including RNA Pol II-mediated regulatory elements (i.e. promoters) (p=9.5 × 10^−4^) and chromatin binding (p=9.5×10^−4^) which is consistent for a multipotent cell type. Among the 31 genes in this category with LoF DNMs, there was a significant enrichment for genes interacting with *SOX2*, a gene recognized to play an essential role in controlling progenitor cell behavior during craniofacial development (P=1.47×10^−3^); the eight genes with LoF DNMs in this category included *MACF1, RBM15, SETD2, CHD7, CTNND1, IRF2BP1, ZFHX4*, and *TFAP2A*. Notably, four of these genes (*CHD7, TFAP2A, ZFHX3*, and *ZFHX4*) were also ranked in the top 100 expressed genes in the lateral nasal eminence (which ultimately forms the upper lip) of E10.5 mouse embryos^29^. Finally, nine of these genes with pLI>0.95 in the top 20^th^ percentile of gene expression had multiple protein-altering DNMs, including *TFAP2A, CTNND1, ZFHX4* and *MACF1*. Ultimately, this analysis provided confirmatory evidence for several genes involved in the etiology of OFCs (*CHD7 and CTNND1)*, expanded the evidence for *TFAP2A* influencing risk to OFCs in humans, and implicated additional genes as novel OFC risk genes (*MACF1, SETD2, ZFHX3*, and *ZFHX4*). To confirm a plausible role in orofacial morphogenesis and cleft pathogenesis, expression patterns of these genes were probed in the embryonic tissues forming the upper lip and primary palate in the mouse. As demonstrated previously, Sox2 expression was localized to the nasal pit epithelium at the center of the lambdoidal junction^30^. All candidate genes examined were detected in the neural crest mesenchyme, with several exhibiting expression adjacent to the Sox2 domain in the nasal pit, including Zfhx4, Mac1, and Setd2.

**Figure 4.**
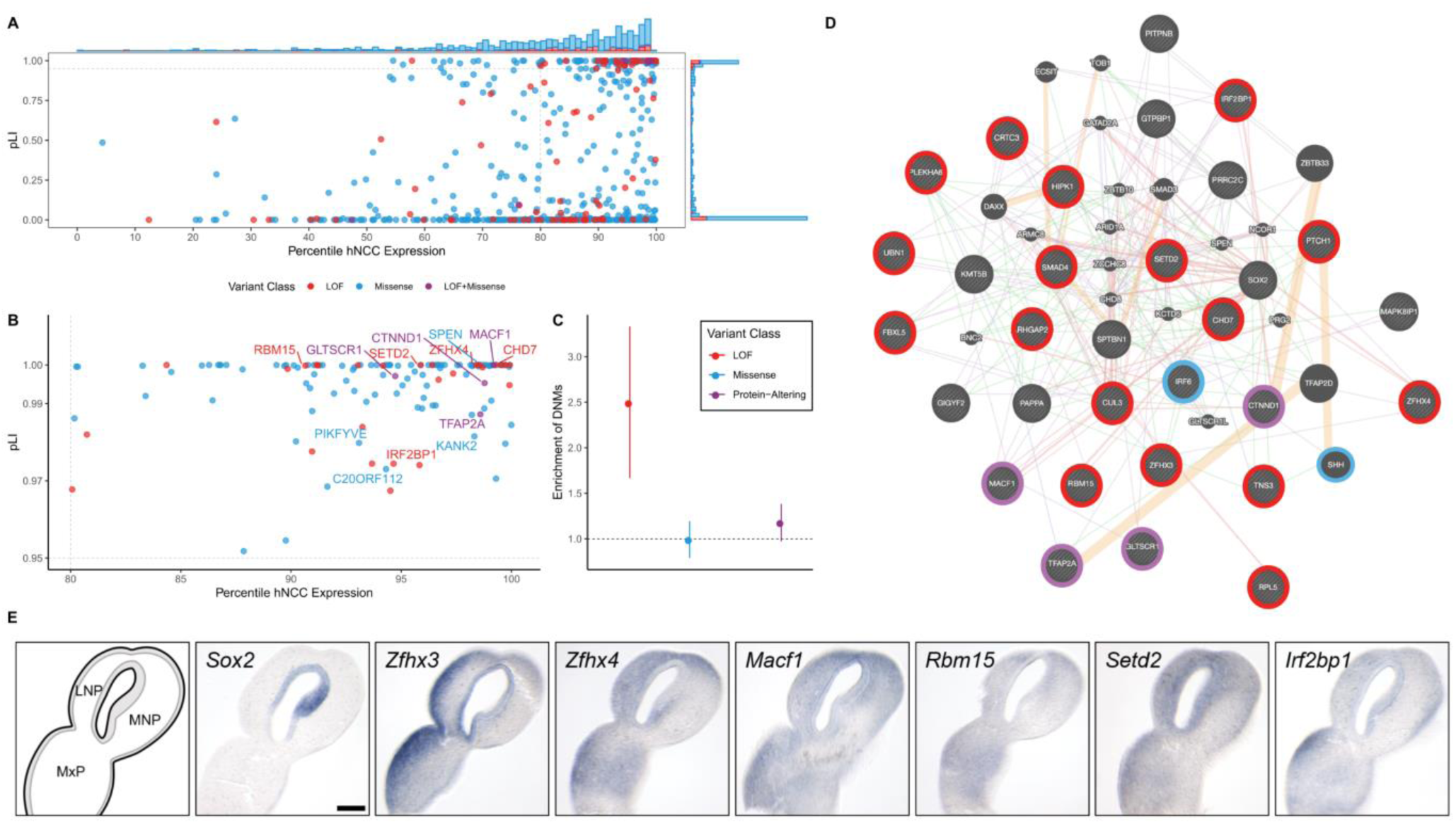
DNMs are enriched in genes expressed in human neural crest cells and among SOX-2 interacting genes. (A) All genes with at least one protein-altering DNM were ranked by their pLI score for the gene and the expression level of the gene in RNA-seq data generated from human neural crest cells. The barplot on the top and right axes show the relative counts of genes with missense (blue), LoF (red), and both missense and LoF (purple) DNMs. (B) Genes with pLI score > 0.95, and in the top 20^th^ percentile of hNCC expression. Labeled genes have at least two protein-altering DNMs. (C) Enrichment of DNMs ± two standard errors in all genes in the top 20^th^ percentile of hNCC expression with a pLI score > 0.95 for all OFC trios. (D) Protein-interaction network generated in GeneMANIA of the 31 genes with LoF DNMs, a pLI score > 0.95, and in the top 20^th^ percentile of hNCC expression. Pink lines represent physical interactions, orange lines represent predicted interactions, purple lines represent co-expression, green lines represent genetic interactions, and blue lines represent co-localization. Genes are colored based on the types of DNMs present in the OFC dataset. (E) Sections through the lambdoidal junction of the medial nasal (MNP), lateral nasal (LNP), and maxillary (MxP) processes of gestational day 11 mouse embryos were stained for the indicated gene by in situ hybridization. In the schematic on the lower left, the epithelium is shaded, while the neural crest mesenchyme is white.

To determine the contribution of DNMs in clinically-relevant OFC genes (Supplementary Table 4), several gene lists were constructed including existing clinical sequencing panels, genes mutated in Mendelian syndromes that include OFCs as a key phenotype, genes previously implicated in candidate gene or exome sequencing studies, and GWAS-nominated genes (see Methods for construction of gene lists). Of the 336 genes comprising this list, we identified 42 individuals with 43 DNMs in 31 genes, representing 6% of all sequenced trios. One individual had two DNMs (a stop-gain in *CHD7* and a synonymous variant in *SHH*), but only the former is predicted to be pathogenic. Overall, 25 missense and 9 loss-of-function DNMs were observed in the list of genes, which was significantly more than expected by chance in this list of genes (P= 8.95 × 10^−6^; Figure 5). This excess was greatest amongst those genes involved in autosomal dominant syndromes (P= 2.29 ×10^−9^). As expected for DNMs, we did not observe an excess among genes involved in autosomal recessive syndromes (P=0.74). GWAS-nominated genes also showed a modest excess of protein-altering DNMs (P= 1.45 × 10^−6^) which in part reflects overlap between these different gene sets (Supplemental Figure 6).

**Figure 5.**
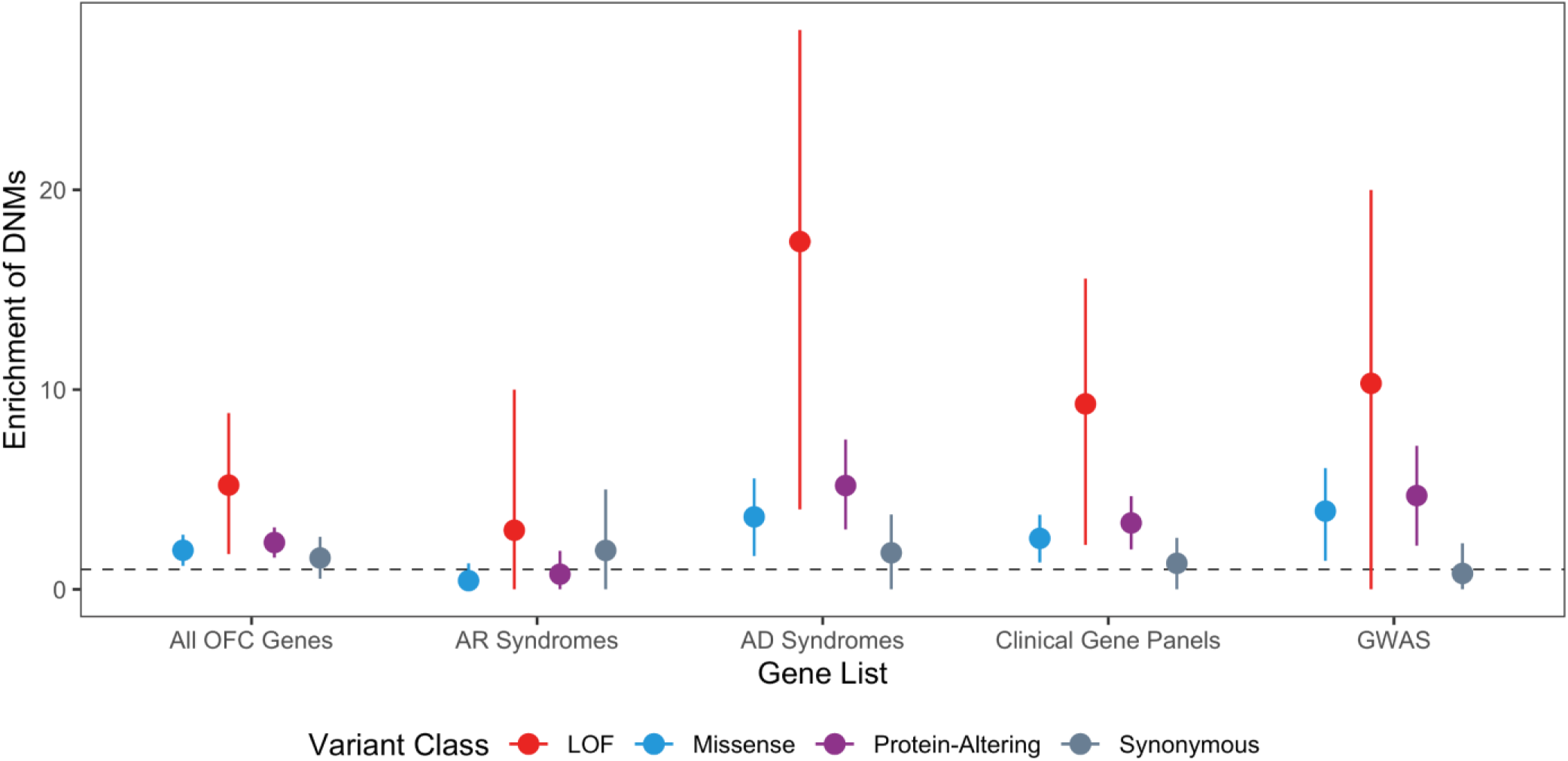
GMKF OFC trios have an 18-fold excess of loss-of-function DNMs in known autosomal dominant OFC genes. Enrichment of DNMs ± two standard errors for all OFC trios in each clinically relevant gene set.

One of the primary challenges in the post-GWAS era is identifying causal genes when the most strongly associated SNPs are non-coding. At some loci, there is an obvious candidate gene where coding mutations can cause Mendelian forms of the same disorder; this is the case for several of the GWAS-nominated genes (e.g., *IRF6, GRHL3, ARHGAP29*). Therefore, we hypothesized identifying a DNM in a gene in the vicinity of a genome-wide association peak could provide evidence to nominate a gene as truly causal or even to provide additional evidence in support suggestive loci not yet achieving genome-wide significance. To address this question, DNMs in genes within 1Mb (± 500kb) of both suggestive and significant GWAS loci from two recent OFC GWASs were evaluated^25; 26^. Overall, 37 protein-altering DNMs were identified within these genes. As expected, several of these DNMs were located within genes already demonstrating strong statistical evidence and implicating them in OFC development. Our results provide confirmatory evidence of some role (*ARHGAP29, IRF6*) in causing OFCs. Protein-altering DNMs were also identified in genes near recently reported GWAS loci (*TFAP2A*^*26*^, *SHROOM3*^*25*^), adding evidence in support of their role in OFC etiology. Furthermore, this analysis also identified genes with DNMs not previously suggested as specific candidate genes within the suggestive or significant loci: *ZFHX4* at 8q21, *RBM15* at 1p13, *UBN1 at 16p13*.*1 and HIRA at 22q11*.*2*, thus providing additional evidence for novel OFC risk genes and loci.

## Discussion

This analysis represents the largest genetic exploration of coding DNMs to date in 756 case-parent trios with non-syndromic OFCs. This initial study clearly demonstrates that coding DNMs in both biologically relevant and clinically relevant sets of genes contribute to the risk of nonsyndromic OFC risk. In addition to providing a unique insight into known OFC risk genes, our results implicate multiple novel candidate genes and gene networks. Collectively, these observations and findings provide a better understanding of the genetic architecture of OFCs.

By combining WGS with single-cell sequencing, bulk RNA-sequencing, and other genomic datasets, we identified multiple novel genes and networks of genes involved in controlling risk to OFCs. The most promising of these genes include *ZFHX3* and *ZFHX4*, the latter of which was the only gene apart from *TFAP2A* with multiple LoF DNMs. Both genes had a pLI score of 1, were highly expressed in the hNCCs(top 20^th^ percentile), and were marker genes for the E0_E11 cell cluster (the olfactory epithelium) analyzed from the single-cell RNA sequencing data. Moreover, *ZFHX4* is located at a suggestive GWAS locus (rs10808812; P=2.00 × 10^−6^)^25^ and is thus implicated in the etiology of OFCs through both LoF DNMs and evidence from common SNPs.

LoF DNMs were also observed in other novel candidates with pLI>0.95 in the top 20^th^ percentile of gene expression, including *MACF1, RBM15*, and *SETD2*. Along with *TFAP2A, CTNND1*, and *CHD7*, all three of these genes interact with *SOX2*, which is known to play an essential role in controlling progenitor cell behavior during craniofacial development^31^. Although *MACF1* was not individually significant in our analyses (P=0.028), we notably identified two protein-altering *MACF1* DNMs (p.Y1357* and p.R5180M). *MACF1* has recently been reported to play a role in regulating osteogenic differentiation and cranial bone formation *in vivo* ^*32*^. *RBM15* is located near a suggestive GWAS locus, providing further evidence for this gene’s involvement in craniofacial pathology. Additionally, mutations in *SETD2* have been reported to cause a novel craniofacial overgrowth condition^33^. Identifying a network of SOX2-interacting genes anchored by established OFC genes paired with the detection of these genes in the neural crest mesenchyme does provide the strongest evidence to support *ZFHX4, ZFHX3, MACF1, RBM15*, and *SETD2* as novel OFC risk genes. Therefore, these genes should be further explored in larger OFC trios.

A major finding of this study is the critical role of *TFAP2A* in OFC risk. There are multiple lines of evidence in the literature to support this assertion, but the number of protein-altering DNMs in *TFAP2A* was unexpected. Complete loss of Tfap2a in mouse models causes severe developmental defects including anencephaly, facial clefts, and thoraco-abdominoschisis^34^. On some backgrounds, heterozygous null mutants exhibit anencephaly^35^. Although the human phenotype caused by *TFAP2A* mutations is considerably less severe^36^, Branchiooculofacial syndrome (BOFS) is described as a very rare disorder with less than 200 described cases as of 2019^37^, corresponding to an estimated prevalence between 1 in 300,000 and 1 in 1,000,000 individuals. BOFS is characterized by problems with branchial arch development leading to skin anomalies on the neck, eye malformations including microphthalmia, anopthalmia, or coloboma, and facial dysmorphism including cleft lip and/or palate, hypertelorism, telecanthus, upslanting palpebral fissures, broad nose, facial muscle weakness, malformed ears, and hearing loss. Given these different phenotypic features, we would expect that the most severely affected BOFS cases would have been excluded from this study so it is unlikely that we would identify 3 cases of classic BOFS in our 756 trios. We conclude that the phenotype caused by mutations in *TFAP2A* mutations is broader than previously appreciated; however, based on our observations, these *TFAP2A* mutations could only account for ∼0.5% of all OFCs.

Approximately 6% of the trios had protein-altering DNMs in clinically-relevant OFC genes, which was significantly more than expected by chance. This finding should be interpreted with caution as this was not a population-based genetic screening of individuals with OFCs. This study population was recruited over many years from multiple sites around the world with a range of clinical skills. Many Mendelian syndromes that include OFCs have variable expressivity, incomplete penetrance, or phenotypic features that may not be apparent at the time of recruitment, especially when cases are often recruited in the first year of life when they present to genetics clinic or begin surgical repairs. Cumulatively, six of the protein-altering DNMs were identified in *IRF6* and *TFAP2A*, accounting for 0.9% of all protein-altering DNMs and 0.8% of sequenced probands. The number of DNMs in IRF6 was expected; dominant mutations in *IRF6* cause Van der Woude syndrome (VWS), the most common Mendelian syndrome with OFC as a key phenotype. VWS cases can mimic nonsyndromic OFCs in about 15% of VWS cases^38^. Other genes with LoF DNMs that are associated with dominant Mendelian syndromes included *RPL5, COL2A1, CTNND1, CHD7*. Recognition of the additional features that characterize syndromes caused by mutations in these genes can become clearer with a molecular finding but it is essential to have a better understanding of how different characteristics of mutations, such as the location within the gene or variant class, contribute to variable expressivity and incomplete penetrance of associated syndromic features. Nonetheless, identifying DNMs in these genes has critical implications for genetic counseling and recurrence risk estimates.

Historically, CL/P and CP have been considered distinct disorders so we performed multiple analyses in CL/P and CP separately to determine if this is supported by DNMs^2^. Both CL/P and CP had a significant excess of protein-altering DNMs, although the excess of DNMs in the CL/P trios was more apparent. The strength of the significance is most likely due to very different sample sizes; 698 trios had CL/P and only 58 had CP. Despite having a small number of CP trios, the GSEA still yielded many significant craniofacial disease terms, although none of these terms was significant for the genes with protein-altering DNMs in the CL/P trios. This lends some support to the idea that CL/P and CP could have distinct etiologies, although we were able to identify genes common to both. Due to these results, the contribution of DNMs should be further investigated in larger studies to better understand causal genetic risk factors and how they contribute differently to OFC subtypes.

The CL/P trios were stratified based on proband sex due to the increased prevalence of CL/P among males compared to females (2:1). In developmental disorders with sex biases, it has been suggested that genetic liability to disease is higher in the less-frequently affected sex and would require more severe variants to manifest the disease (i.e. a higher threshold determining affected vs. not in the sex with lower population prevalence). For OFCs, we would hypothesize that females with CL/P would have an increased burden of DNMs compared to males or that they would have more LoF DNMs. However, males and females did not differ significantly by any DNM variant class and both sexes had a similar significant excess of LoF and protein-altering DNMs. These results suggest that the difference in genetic liability cannot be explained by coding, autosomal DNMs alone.

In conclusion, we have shown the important contribution of rare, coding DNMs in 756 OFC case-parent trios. Through this exploration we have identified novel risk genes as well as pertinent information regarding well-known and clinically-relevant OFC genes. Like other structural birth defects, both missense and LoF DNMs play a significant role in determining risk to OFC. Similar studies in specific OFC subtypes will be necessary to fully understand the genetic architectures specific to each cleft subtype separately. Our results also suggest that the analysis of rare inherited variants in these same genes will uncover additional risk variants and may identify some of the missing heritability for OFCs. More challenging will be to explore the implications of noncoding variants, but this study highlights sets of genes and pathways that should be the focus for identifying possible regulatory elements.

## Supplementary Material

Supplementary material contains 7 figures and 5 tables.

## Acknowledgements

These studies are part of the Gabriella Miller Kids First Pediatric Research Program consortium, supported by the Common Fund of the Office of the Director of the National Institutes of Health (www.commonfund.nih.gov/KidsFirst). Sequencing of the European trios was completed at Washington University’s McDonnell Genome Institute (3U54HG003079-12S1, X01-HL132363 [MLM, EF]) and the Colombian and Taiwanese trios were sequenced at the Broad Institute Sequencing Center (U24-HD090743, X01-HL136465 [MLM, EF], X01-HL140516 [THB, Azeez Butali]). The sequencing centers plus the Kids First Data Resource Center (kidsfirstdrc.org, supported by the NIH Common Fund through U2CHL138346) provided technical and analytical support of this project. The data analyzed and reported in this manuscript were accessed from dbGaP [www.ncbi.nlm.nih.gov/gap; European trios: dbGaP accession number phs001168.v2.p2; Colombian trios: dbGaP accession number phs001420.v1.p1; Taiwanese trios: phs000094.v1.p1] and from the Kids First Data Resource Center (kidsfirstdrc.org). Additional grants supported the assembling of the sample of case-parent trios, collection of the phenotypic data and samples, and data analysis were supported by the NIH: R01-DE016148 [MLM, SMW], R03-DE026469 [EF, MLM], R03-DE027193 [EJL], R00-DE025060 [EJL], R01-DE011931 [JTH], U01-DD000295 [GW], R03-DE027121 [MAT, THB, JB]. This study would not be possible without the dedication of many families, study teams, and colleagues worldwide.

## Author Contributions

Conceptualization, E.J.L; Methodology, M.R.B., H.B., M.P.E., M.R.S., R.J.L., D.J.C, E.J.L; Software, M.R.B., P.C., N.M., D.J.C.; Formal Analysis, M.R.B., P.C., D.J.C., M.R.S., R.J.L., I.R., E.J.L; Investigation, M.R.B., K.D.P, M.R.S., S.H.; Data Curation, M.R.B., N.M., J.H., M.A.T.; Writing-Original Draft, M.R.B., E.J.L; Writing-Review and Editing, All Authors; Visualization, M.R.B., E.J.L; Supervision, E.J.L, E.F., D.J.C, M.L.M; Resources, L.M.M-U, L.C.V-R, C.R., G.W., J.T.H, F.D., S.M.W, Y.H.W-C, P.K.C, T.H.B, J.C.M, M.L.M, E.J.L; Project Administration, E.J.L; Funding Acquisition, E.J.L, E.F., M.L.M, S.M.W, T.H.B.

