## Supplemental for "Coding *de novo* mutations identified by WGS reveal novel orofacial cleft genes"

### Supplementary Material

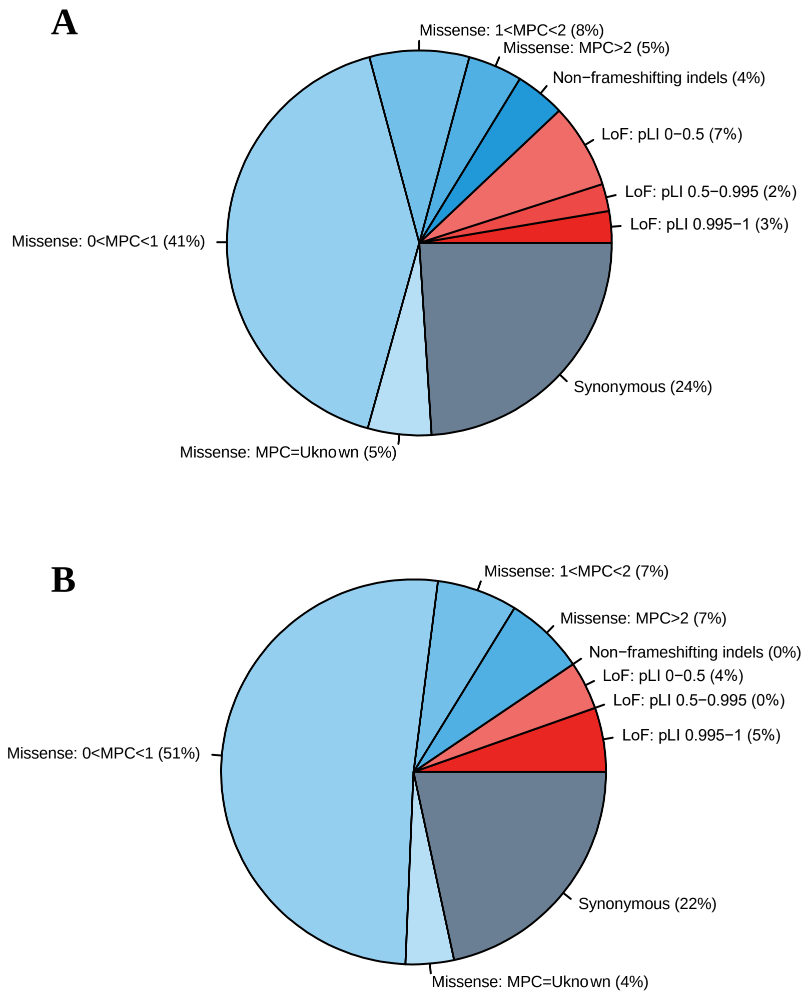


**Supplementary Figure 1**. **(A)** Distribution of rare, coding DNMs by variant class for CL/P trios. **(B)** Distribution of rare, coding DNMs by variant class for CP only trios.
DNMs were subcategorized by MPC score (missense) or pLI score (LoF).

A

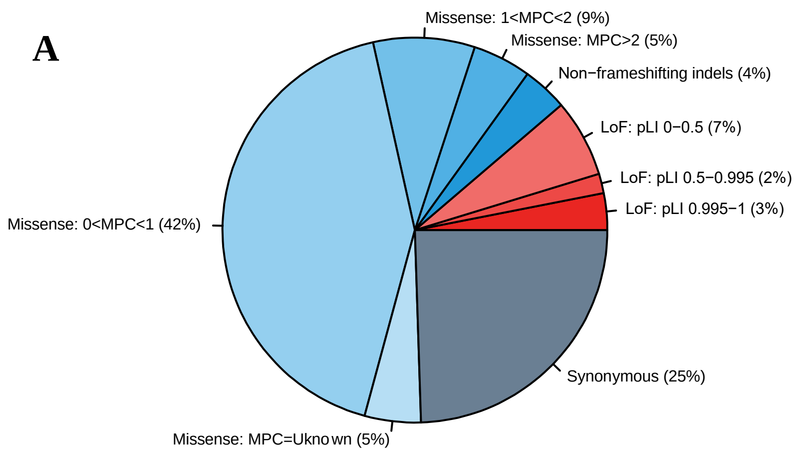


B

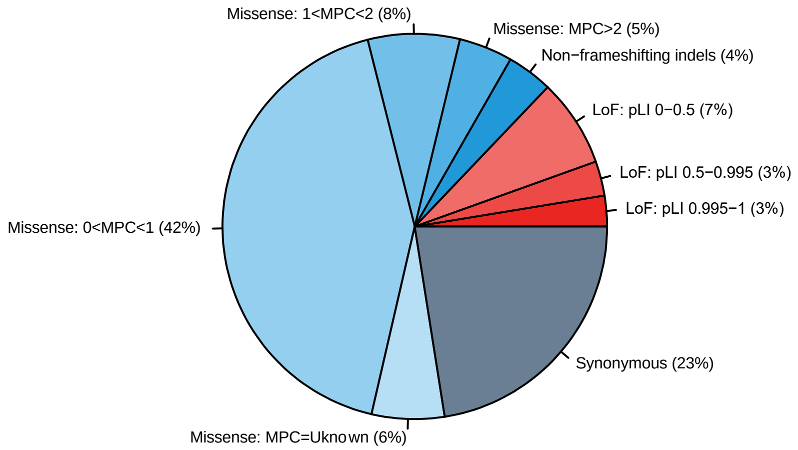


**Supplementary Figure 2**. **(A)** Distribution of rare, coding DNMs by variant class for male probands with any OFC. **(B)** Distribution of rare, coding DNMs by variant class for female probands with any OFC.
DNMs were subcategorized by MPC score (missense) or pLI score (LoF)

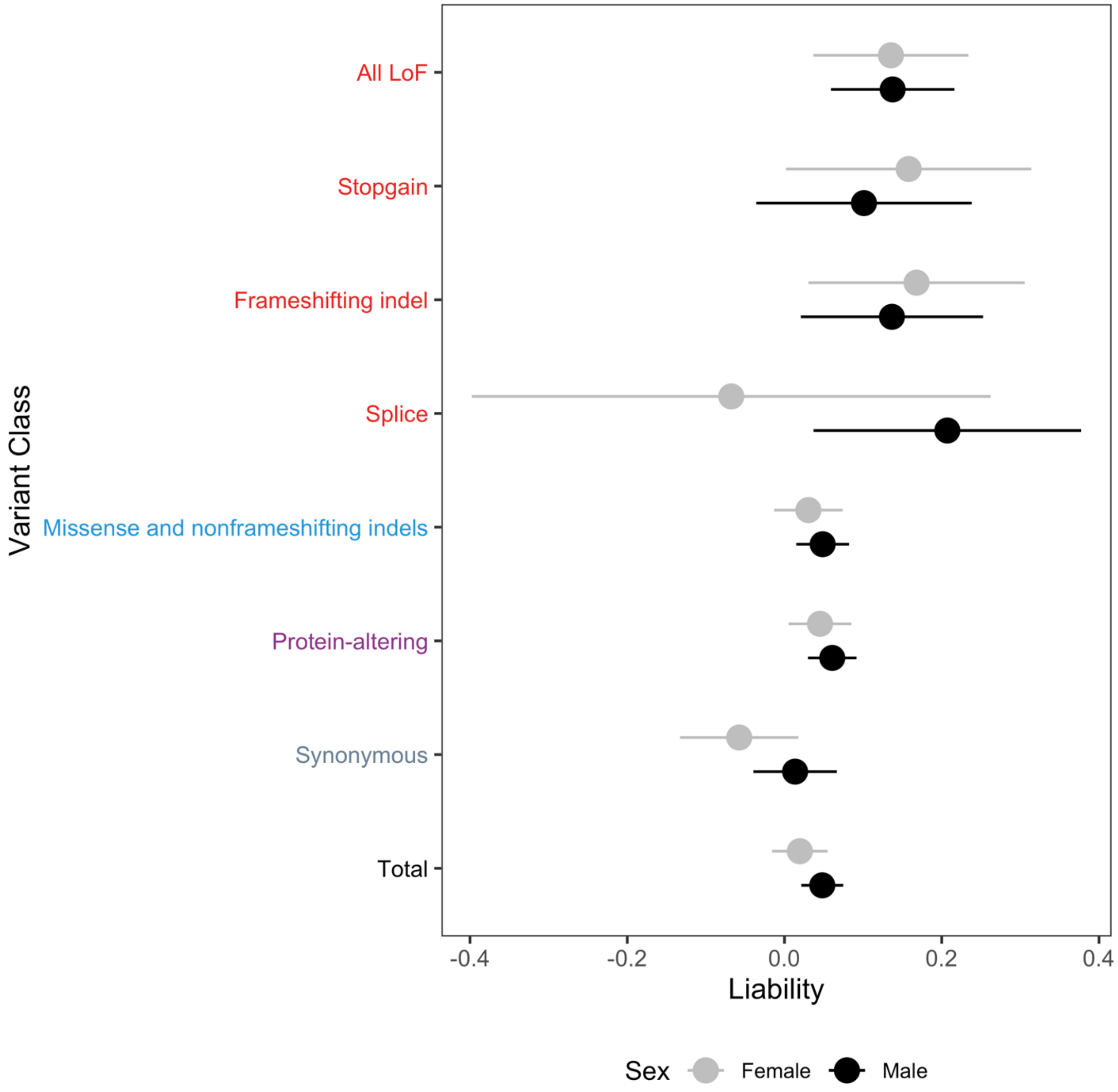


**Supplementary Figure 3.** Comparison of the number of DNMs by variant type between males and females on the liability scale.

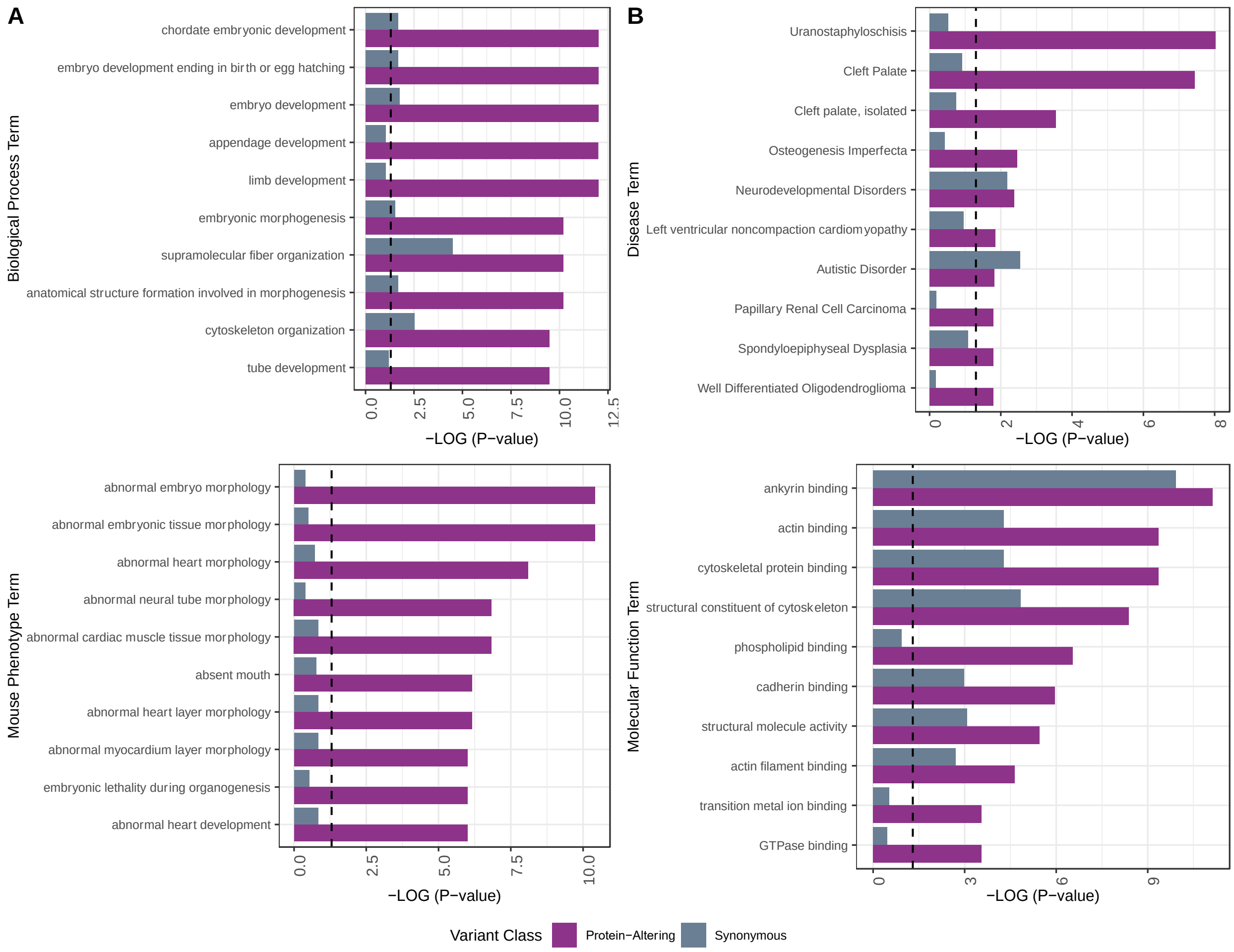


**Supplementary Figure 4.** Gene set enrichment analysis for all OFC for genes with protein-altering (purple) or synonymous (grey) DNMs. The dashed line represents a significance threshold of p-value=0.05. **(A)** P-values for genes with synonymous DNMs and protein-altering DNMs for the top ten most significant biological process terms (top) and mouse phenotype terms (bottom) for genes with protein-altering DNMs. **(B)** P-values for genes with synonymous DNMs and protein-altering DNMs for the top ten most significant disease terms (top) and molecular function terms (bottom) for genes with protein-altering DNMs.

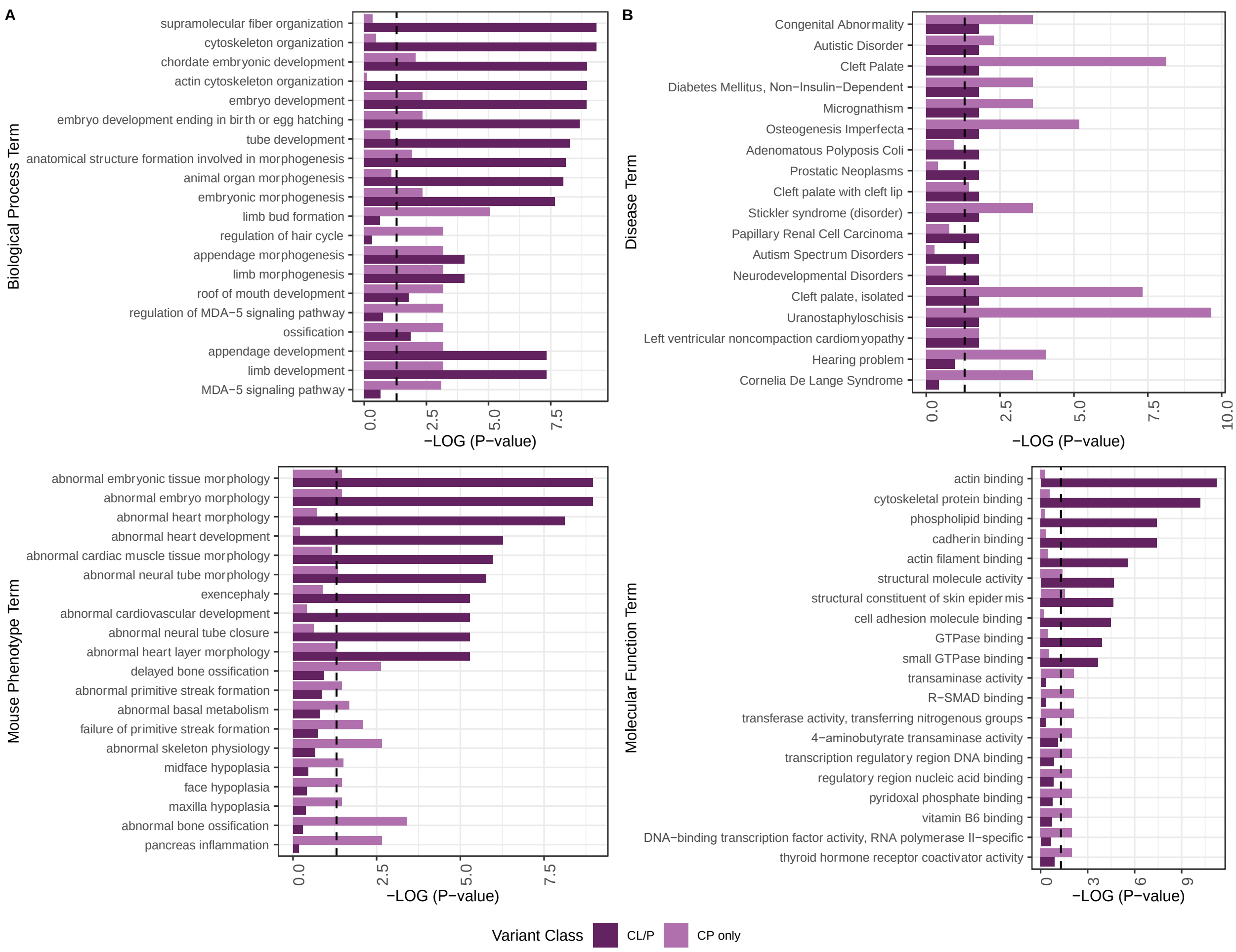


**Supplementary Figure 5.** Gene set enrichment analysis for genes with protein-altering DNMs in the CL/P trios (dark purple) and the CP only trios (light purple). **(A)** P-values for the top ten most significant biological process terms (top) and mouse phenotype terms (bottom) for genes with protein-altering DNMs in CL/P and CP only trios. **(B)** P-values for the top ten most significant disease terms (top) and molecular function terms (bottom) for genes with protein-altering DNMs in CL/P and CP only trios.

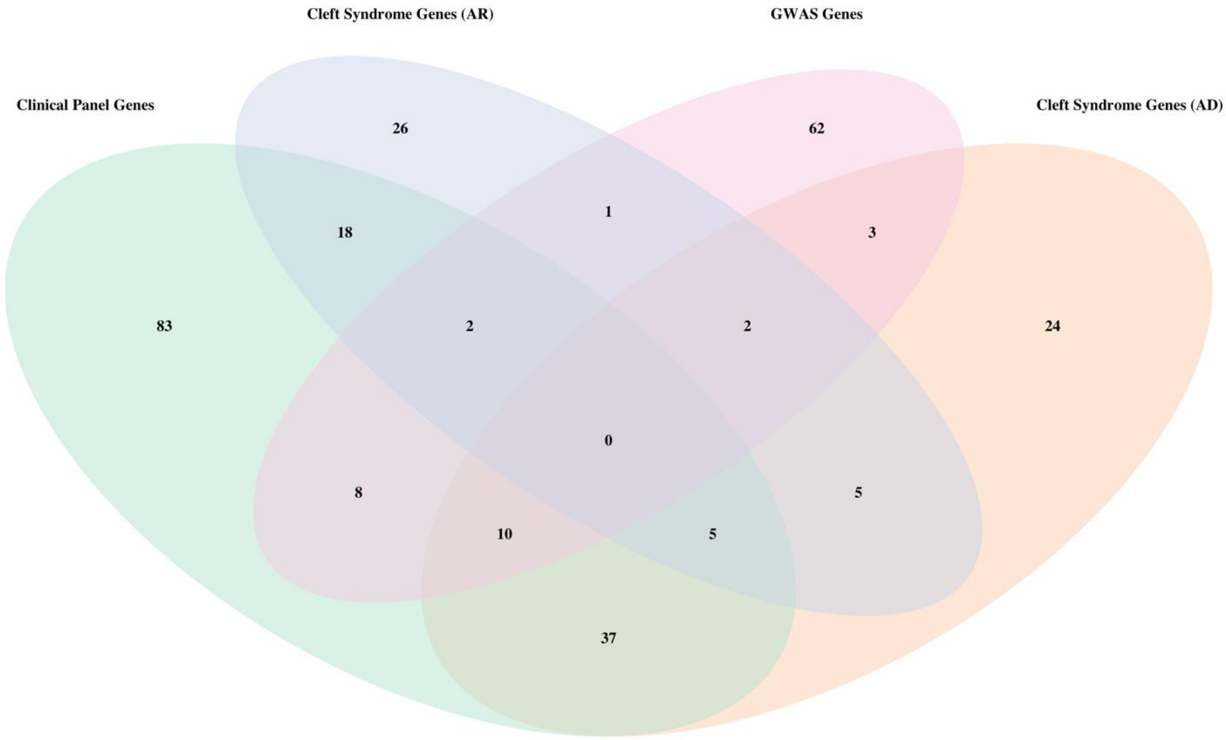


**Supplementary Figure 6.** Venn diagram showing the number of genes in each clinically relevant gene set list.

A

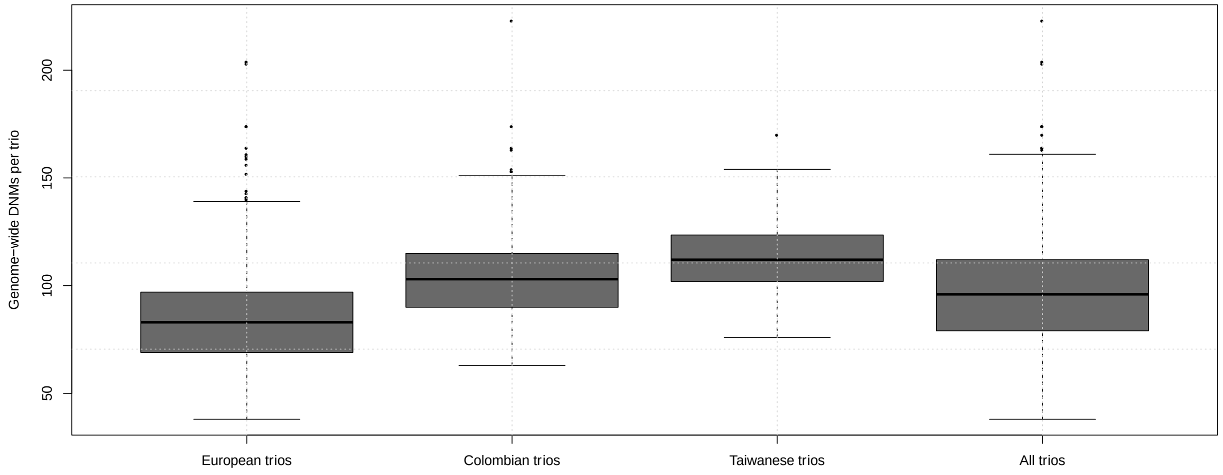


B

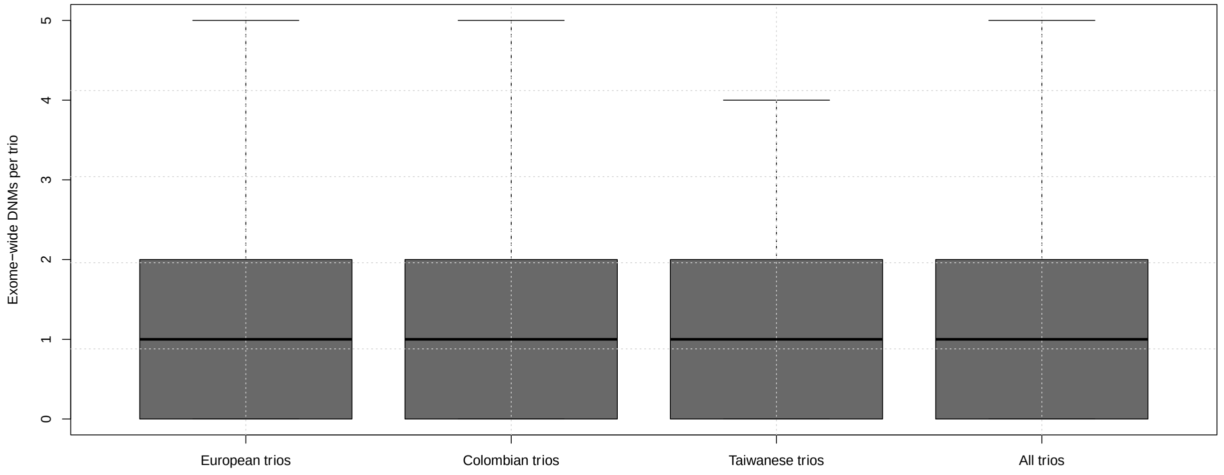


**Supplementary Figure 7.** Comparison of the number of DNMs per trio by ethnicity. (**A)** Genome-wide DNMs per trio for European, Colombian, Taiwanese, and All trios. **(B)** Exome-wide DNMs per trio for European, Colombian, Taiwanese, and All trios.

**Supplementary Table 1.** Summary of the GMKF sample of case-parent trios with OFCs

| Sample | Total Trios | Trios with no affected parents | Trios with 1 affected parent | Trios with 2 affected parents | Offspring cleft status | | | | | |
|---|---|---|---|---|---|---|---|---|---|---|
| | | | | | CL/P | | | CP only | | |
| | | | | | Total | Male | Female | Total | Male | Female |
| **European** | 373 | 331 | 38 | 4 | 315 | 209 | 106 | 58 | 32 | 26 |
| **Colombian** | 267 | 267 | 0 | 0 | 267 | 156 | 111 | 0 | 0 | 0 |
| **Taiwanese** | 116 | 108 | 8 | 0 | 116 | 79 | 37 | 0 | 0 | 0 |
| **Total** | 756 | 706 | 46 | 4 | 698 | 444 | 254 | 58 | 32 | 26 |

**Supplementary Table 2.** Summary of DNMs identified the GMKF sample of case-parent trios with OFCs

| Variant Class | Variant Class Subclassification | Colombian CL/P (N) | | | European CL/P (N) | | | Euro. CP (N) | Taiwanese CL/P (N) | | | All CL/P (N) | | | All OFC (N) | | |
|---|---|---|---|---|---|---|---|---|---|---|---|---|---|---|---|---|---|
| | | All (267) | M (156) | F (111) | All (315) | M (210) | F (105) | All (58) | All (116) | M (79) | F (37) | All (698) | M (445) | F (253) | All (756) | M (477) | F (279) |
| **Loss of Function** | Total | 41 | 25 | 16 | 42 | 25 | 17 | 7 | 12 | 10 | 2 | 95 | 60 | 35 | 102 | 62 | 40 |
| | Stopgain | 15 | 8 | 7 | 17 | 10 | 7 | 4 | 2 | 2 | 0 | 34 | 20 | 14 | 38 | 21 | 17 |
| | Frameshifting indel | 18 | 10 | 8 | 18 | 10 | 8 | 1 | 9 | 7 | 2 | 45 | 27 | 18 | 46 | 27 | 19 |
| | Splice | 8 | 7 | 1 | 7 | 5 | 2 | 2 | 1 | 1 | 0 | 16 | 13 | 3 | 18 | 14 | 4 |
| | Loss of Function pLI:0.995-1 | 10 | 9 | 1 | 8 | 3 | 5 | 4 | 3 | 3 | 0 | 21 | 15 | 6 | 25 | 17 | 8 |
| | Loss of Function pLI: 0.5-0.995 | 11 | 4 | 7 | 5 | 3 | 2 | 0 | 2 | 2 | 0 | 18 | 9 | 9 | 18 | 9 | 9 |
| | Loss of Function pLI: 0-0.5 | 20 | 12 | 8 | 29 | 19 | 10 | 3 | 7 | 5 | 2 | 56 | 36 | 20 | 59 | 36 | 23 |
| **Non-frameshifting indels** | | 12 | 9 | 3 | 12 | 8 | 4 | 0 | 9 | 4 | 5 | 33 | 21 | 12 | 33 | 21 | 12 |
| **Missense** | Total | 201 | 118 | 83 | 194 | 133 | 61 | 51 | 76 | 55 | 21 | 471 | 306 | 165 | 522 | 333 | 189 |
| | MPC>2 | 15 | 8 | 7 | 12 | 9 | 3 | 5 | 9 | 6 | 3 | 36 | 23 | 13 | 41 | 27 | 14 |
| | MPC: 1-2 | 20 | 13 | 7 | 36 | 23 | 13 | 5 | 10 | 7 | 3 | 66 | 43 | 23 | 71 | 47 | 24 |
| | MPC:0-1 | 148 | 88 | 60 | 129 | 90 | 39 | 38 | 50 | 38 | 12 | 327 | 216 | 111 | 365 | 233 | 132 |
| | Unknown | 18 | 9 | 9 | 17 | 11 | 6 | 3 | 7 | 4 | 3 | 42 | 24 | 18 | 45 | 26 | 19 |
| **Synonymous** | | 76 | 49 | 27 | 75 | 56 | 19 | 16 | 38 | 26 | 12 | 189 | 131 | 58 | 205 | 135 | 70 |
| **Protein-altering** | | 254 | 152 | 102 | 248 | 166 | 82 | 58 | 97 | 69 | 28 | 599 | 387 | 212 | 657 | 416 | 241 |
| **Total** | | 330 | 201 | 129 | 323 | 222 | 101 | 74 | 135 | 95 | 40 | 788 | 518 | 270 | 862 | 551 | 311 |

**Supplementary Table 5.** Summary of gene-specific ISH riboprobe primers used for in situ hybridization

| **Species** | **Primer** | **Sequence** |
|---|---|---|
| Mouse | Irf2bp1 F | GCTTCAAGTACCTCGAGTATG |
| Mouse | Irf2bp1 R | <u>CGATGTTAATACGACTCACTATAGGG</u> TGATGTCACCAGCAAGAATAG |
| Mouse | Macf1 F | CTTACAACAGGAGACAGAGAAG |
| Mouse | Macf1 R | <u>CGATGTTAATACGACTCACTATAGGG</u> TAGAGTGGAGAGTGGTGTATC |
| Mouse | Rbm15 F | AACGCTTCGGTGATGTAAG |
| Mouse | Rbm15 R | <u>CGATGTTAATACGACTCACTATAGGG</u> GGCCTCTTAATGTCCACTTC |
| Mouse | Setd2 F | AGTCCTCCGTCAGGAATAAG |
| Mouse | Setd2 R | <u>CGATGTTAATACGACTCACTATAGGG</u> GGAGTCGGTTTCTTGGAATAC |
| Mouse | Sox2 F | GAAGGATAAGTACACGCTTCC |
| Mouse | Sox2 R | <u>CGATGTTAATACGACTCACTATAGGG</u> GCGTTAATTTGGATGGGATTG |
| Mouse | Zfhx3 F | ACAGCGCAACAGGAATAG |
| Mouse | Zfhx3 R | <u>CGATGTTAATACGACTCACTATAGGG</u> GATACGTGGTAGGAAGGTTAAG |
| Mouse | Zfhx4 F | CTTGACCGGGAGAAAGATTAC |
| Mouse | Zfhx4 R | <u>CGATGTTAATACGACTCACTATAGGG</u> GTTTGATAGCCTCCGATTCC |
